# The dorsal and ventral default mode networks are dissociably modulated by the valence and vividness of imagined events

**DOI:** 10.1101/2020.05.21.109728

**Authors:** Sangil Lee, Trishala Parthasarathi, Joseph W. Kable

**Affiliations:** Department of Psychology, University of Pennsylvania, Philadelphia, PA, 19104. USA; Department of Neuroscience, University of Pennsylvania, Philadelphia, PA, 19104. USA

## Abstract

Recent work has shown that the brain’s default mode network (DMN) is active when people imagine the future. Here we test whether future imagination can be decomposed into two dissociable psychological processes linked to different subcomponents of the DMN. While measuring brain activity with fMRI as subjects imagine future events, we manipulate the vividness of these events to modulate the demands for scene construction, and we manipulate the valence of these events to modulate the demands for evaluation. We found that one subcomponent of the DMN, the ventral DMN or medial temporal lobe subsystem, responds to the vividness but not the valence of imagined events. In contrast, another subcomponent, the dorsal or core DMN, responds to the valence but not the vividness of imagined events. This separate modifiability of different subcomponents of the DMN by vividness and valence provides strong evidence for a neurocognitive dissociation between (1) the construction of novel, imagined scenes from individual components from memory and (2) the evaluation of these constructed events as desirable or undesirable.

**Significance Statement:** Previous work has suggested that imagination may depend on separate neural networks involved in the construction and evaluation of imagined future events. This study provides strong neural evidence for this dissociation by demonstrating that two components of the brain’s default mode network (DMN) uniquely and specifically respond to different aspects of imagination. The vividness of imagined events modulates the ventral DMN, but not the dorsal DMN, while the valence of imagined events modulates the dorsal DMN, but not the ventral DMN. This supports the dissociable engagement of these sub-networks in constructing and evaluating imagined future events.

## Introduction

Imagining the future can aid planning and help one act advantageously in the present. But what is the underlying cognitive architecture of imagination? Though imagination, like perception, can subjectively feel like a unitary experience, it may arise from the interaction of dissociable psychological processes. Here we investigate the hypothesis that imagination consists of at least two distinct processes: a *constructive* process, by which a novel future event is mentally formed, often by combining specific aspects of past experience (Addis, Wong, & Schacter, 2007; Hassabis, Kumaran, & Maguire, 2007; Schacter, Addis, & Buckner, 2007); and an *evaluative* process, by which the imagined event is judged as positive or negative (D’Argembeau & Van Der Linden, 2004; Gilbert & Wilson, 2007; Sharot, Riccardi, Raio, & Phelps, 2007).

Because imagination is fundamentally an internal, subjective activity, studying its architecture can be difficult with behavioral data alone, and therefore many studies have turned to brain imaging. These studies have often focused on the default mode network (DMN), as envisioning and evaluating future events is proposed to be a key function of the DMN (Addis et al., 2007; Botzung, Denkova, & Manning, 2008; Okuda et al., 2003; Sharot et al., 2007; Szpunar, Watson, & McDermott, 2007). DMN is one of the core networks reliably recovered from resting-state fMRI studies and includes the ventromedial prefrontal cortex (vmPFC), posterior cingulate cortex (PCC), and regions in the medial temporal and parietal lobes, such as hippocampus and precuneus (Andrews-Hanna, Reidler, Sepulcre, Poulin, & Buckner, 2010; Andrews-Hanna et al., 2007; Greicius, Srivastava, Reiss, & Menon, 2004; Raichle, 2015; Spreng, Mar, & Kim, 2009).

Past research suggests that constructive and evaluative processes may engage different components of the DMN. Studies of “scene construction,” when elements of the past are combined to create a novel potential future event, have revealed activity in the hippocampus, parahippocampal gyrus, and retrosplenial cortex (Addis & Schacter, 2008; Andrews-Hanna et al., 2007; Greicius et al., 2004; Hassabis et al., 2007; Hassabis & Maguire, 2007). In contrast, activity in vmPFC is seen in tasks with an evaluative component. Activity in vmPFC is associated with the value of predicted future outcomes, and imagining positive events increases activity in vmPFC compared to imagining negative or control events (Bartra, McGuire, & Kable, 2013; D’Argembeau, 2013; D’Argembeau, Xue, Lu, Van der Linden, & Bechara, 2008; Roy, Shohamy, & Wager, 2012; Sharot et al., 2007).

Resting-state functional connectivity studies also point to sub-divisions of the DMN. Using seed-based resting-state functional connectivity, Andrews-Hanna et al. (2010, 2017) distinguished between a medial temporal lobe (MTL) subsystem, consisting of hippocampal, parahippocampal, restrosplenial, ventromedial prefrontal, and posterior parietal cortex, and a DMN core, consisting of medial prefrontal cortex and posterior cingulate cortex. Using independent components analysis, Shirer et al. (2012) proposed a similar distinction between a dorsal DMN, which largely overlaps with the DMN core, and a ventral DMN, which largely overlaps with the MTL subsystem.

Here we constructed a strong test of the hypothesis that the DMN consists of dissociable constructive and evaluative networks involved in future imagination. We relied on the logic of *separate modifiability* (Sternberg, 2001), which supports stronger inferences regarding dissociations, by showing that two processes are differentially influenced by distinct factors within the same task. To modulate activity in brain regions engaged in constructive processes during imagination, we manipulated the vividness of imagined events, where vividness refers to the amount of detail or concreteness of the imagined event. To modulate activity in brain regions engaged in evaluative processes during imagination, we manipulated the valence of imagined events, where valence refers to the intensity of positive or negative emotions the imagined event invokes. If one component of the DMN is modulated by the vividness but not the valence of imagined events, while another component is modulated by valence but not vividness, this double dissociation would provide strong evidence for a functional division of the DMN associated with constructive versus evaluative processes.

## Methods

### Subjects

Twenty-four participants (13 females, average age = 24.9 years, SD = 4.6 years) were recruited from the University of Pennsylvania and surrounding community. One additional participant was excluded for excessive head movement (shifts of at least 0.5 mm between >5% of adjacent time points). All participants were compensated for their time at $15 per hour and provided consent prior to study procedures in accordance with the procedures of the Institutional Review Board of the University of Pennsylvania.

### Imagination task

All participants completed an imagination task in the scanner. Participants were asked to imagine scenarios and then rate the imagined scenarios on vividness and valence (Figure 1). Thirty-two scenarios were presented in each run and participants completed a total of 4 runs. The vividness and valence ratings were performed on a 7-point Likert Scale. To assess vividness, participants were asked “How vividly did you imagine this event” with anchors of “Vague with no details” to “Vividly clear.” To assess valence, participants were asked “How would you rate the valence of emotions in this event” with anchors of “Very Negative” to “Very Positive.”

**Figure 1.**
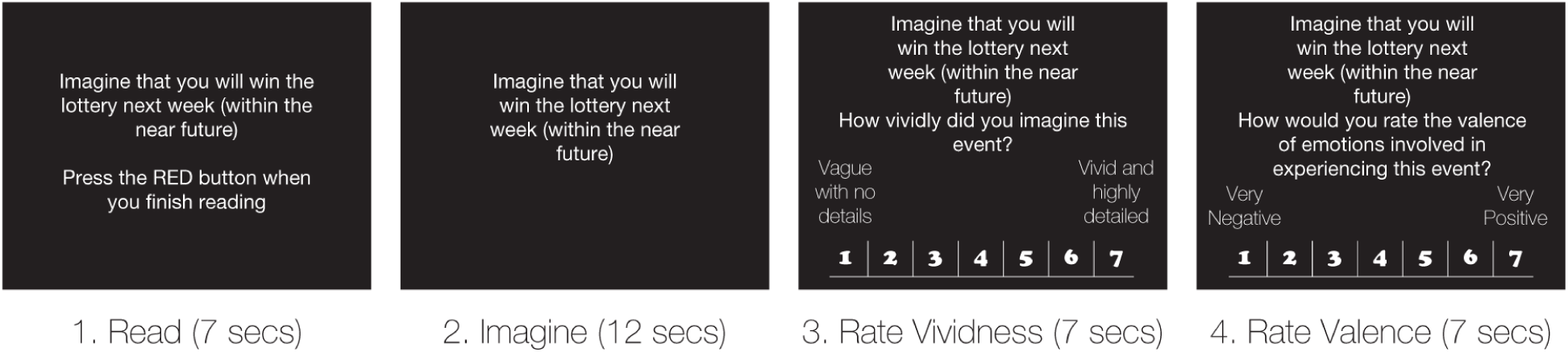
Experimental procedure of the task. Participants had up to 7 secs to read the cue, 12 seconds to imagine, and up to 7 seconds each to rate the vividness and valence of the scenario.

Participants were given up to 7 seconds to read the cue, 12 seconds to imagine the scenario, up to 7 seconds to rate vividness, and up to 7 seconds to rate valence. The participant pressed a button indicating that the cue was read to start the imagination epoch. The imagination epoch was a fixed 12 seconds for all participants. Following imagination, participants were given up to 7 seconds to move a scale ranging from 1-7 to make their rating. If participants failed to submit a rating response within the allotted time, the last rating the participant had highlighted at that point was taken as their selection. Any time not used in any of the free response intervals was added to the inter-trial-interval, so that a new trial occurred every 33 seconds.

### Scenarios

The scenarios were selected for high or low vividness and for positive or negative valence. A list of 68 distinct scenarios was compiled from other studies that assessed vividness, valence, and other aspects of imagination, as well as a survey of MTURK respondents (n = 411, 199 female, average age = 30.1 years, SD = 11 years) who were given broad categories of possible scenarios and asked to create their own. These scenarios were then rated in a separate study (n = 131, 73 female, average age = 34.6 years, SD = 12 years) on MTURK, with each participant rating the valence and vividness of 17 of the 68 scenarios. Based on these ratings, a final list of 32 scenarios was created by selecting the most and least vivid positive and negative scenarios. The final stimulus set included 8 scenarios in each of four conditions – Vivid Positive, Vivid Negative, Non-Vivid Positive, and Non-Vivid Negative.

To more exhaustively characterize the differences between vivid and non-vivid and positive and negative scenarios, we performed a further online survey (n = 391, average age = 37.3 years, SD = 11.8 years). Online participants read each scenario and answered a question about the imagined event. A different group of 32 participants answered for each of the following 12 different measures:

1. Arousal: What was your level of arousal in experiencing this event? (1 = Not at all, 7 = Extremely)
2. Current Emotion: How intense is the emotion felt at the time of imagining the event? (1 = Not at all intense, 7 = Extremely intense)
3. Future Emotion: How intense would your emotion be at the time when the future event takes place? (1 = not at all intense, 7 = extremely intense)
4. Personal Importance: What is the personal importance of this event? (1 = not important, 7 = extremely important)
5. Pre-Experience: How much did you pre-experience the imagined event? (How much did you feel like you were actually there?) (1 = Not at all, 7 = completely)
6. Self-Relevance: How relevant is the imagined event to you? (1 = not at all relevant, 7 = extremely relevant)
7. Social Connection: How much did imagining this event make you feel connected to other people? (1 = not at all connected, 7 = very connected)
8. Subjective Temporal Distance: How far away do you feel from the imagined future event? (1 = very close, 7 = very far)
9. Temporal Connection: What is the perceived similarity of your current self to your self in the imagined future event? (1 = very different, 7 = exactly the same)
10. Visual Perspective: What is your perspective when imagining this event? Are you actively participating (field) or simply observing (observer)? (1 = field, 7 = observer)
11. Valence: How would you rate the valence of emotions involved in experiencing this event? (1 = very negative, 7 = very positive)
12. Vividness: How vividly did you imagine this event? (1 = vague with no details, 7 = vivid and highly detailed)

The biggest difference between vivid and non-vivid scenarios was in vividness, but consistent with vividness affecting constructive processes, people were more likely to imagined vivid scenarios as active participants rather than observers (Table 1). The biggest difference between positive and negative scenarios was in valence, but consistent with valence affecting evaluative processes, people reported more arousal, less emotional intensity, a greater sense of social and temporal connectedness and more self-relevance for positive scenarios (Table 1). Both vivid and positive scenarios were associated with a greater feeling of being “actually there.”

**Table 1.** Further characterization of vivid versus non-vivid and positive versus negative scenarios in separate online sample. Average ratings for each of fourteen different questions are provided and compared across non-vivid vs. vivid scenarios and positive vs. negative scenarios. Ratings were made of arousal, current emotion, future emotion, personal importance, pre-experience, self-relevance, social connection, subjective temporal distance, temporal connection, visual perspective, valence, and vividness.

|  | Arousal | Curr.<br>Emot. | Fut.<br>Emot. | Pers.<br>Imp. | Pre-Exp | Self-Rel | Soc.<br>Conn. | Temp.<br>Dist. | Temp.<br>Conn. | Vis.<br>Pers. | Valence | Vivid |
| --- | --- | --- | --- | --- | --- | --- | --- | --- | --- | --- | --- | --- |
| Nonvivid | 4.56 | 4.43 | 5.04 | 5.27 | 4.86 | 4.40 | 3.96 | 4.33 | 4.31 | 3.82 | 3.90 | 4.56 |
| Vivid | 4.86 | 4.73 | 5.26 | 4.92 | 5.35 | 4.89 | 3.88 | 3.89 | 4.75 | 3.26 | 4.06 | 5.34 |
| *t*-test *p* | .153 | .242 | .374 | .267 | .002* | .028 | .871 | .218 | .134 | .001* | .823 | <.001* |
| Negative | 4.42 | 4.92 | 5.41 | 5.02 | 4.83 | 4.35 | 3.01 | 4.45 | 4.06 | 3.62 | 2.11 | 4.86 |
| Positive | 4.99 | 4.25 | 4.89 | 5.16 | 5.38 | 4.93 | 4.83 | 3.77 | 4.99 | 3.46 | 5.85 | 5.04 |
| *t*-test *p* | .004* | .007* | .032* | .663 | .001* | .010* | <.001* | .051 | .001* | .356 | <.001* | .364 |

In the scanner, these 32 scenarios were repeated with both “near future” and “far future” prompts, where “near future” was defined as “within the next week” and “far future” was defined as “more than a year from now”. This resulted in 64 unique scenario prompts, with each run containing 16 prompts. For the scanner participants, we conducted a repeated-measures ANOVA to ensure that there was a main effect of valence and vividness of the scenario on participants’ judgments of valence and vividness.

### Imaging Acquisition

Functional and anatomical images were collected using a 3T Siemens Trio scanner equipped with a 32-channel head coil. At the beginning of each session, high-resolution T1-weighted anatomical images were collected using an MPRAGE sequence (T1 = 1100ms; 160 axial slices, 0.9375 × 0.9375 × 1.000 mm; 192 × 256 matrix). T2*-weighted functional images were then collected using an EPI sequence (3 mm isotropic voxels, 64 × 64 matrix, 44 axial slices tilted 30o from the AC-PC plane, TR = 3,000ms, TE = 25ms). All participants completed four functional scans in each session, with each functional scan consisting of 181 images. At the end of the session we acquired matched B0 fieldmap images (TR = 1000ms, TE = 2.69 and 5.27ms). These images as well as the behavioral responses are available online at openneuro.org (DOI: 10.18112/openneuro.ds002835.v1.0.0).

### Imaging Analyses and Preprocessing

Brain imaging analysis was conducted with the FMRIB Software Library (FSL) using FSL FEAT (FMRIB fMRI Expert Analysis Tool) version 6.00 (Smith et al., 2004). Preprocessing included the following: (1) skull stripping of structural images with BET (FMRIB Brain Extract Tool); (2) motion correcting with MCFLIRT (FMRIB Linear Image Restoration Tool with Motion Correction); (3) spatial smoothing with a 9mm full-width half-maximum Gaussian kernel; and (4) high-pass temporal filtering equivalent to 150 Hz. Registration and normalization were performed with FLIRT. Each functional image was registered to the participant’s high-resolution brain-extracted structural image using boundary-based registration that simultaneously incorporates fieldmap-based geometric distortion and normalized to the FSL Montreal Neurological Institute (MNI) template using affine transformations with 12 degrees of freedom.

We focused on testing which aspects of the imagined events significantly modulated neural activity in different regions of the brain. Our general linear model included regressors for the Read, Imagine, Rate Vividness, and Rate Valence epochs, as well as categorical event modulators for the Imagine epoch for the Vividness (high versus low), Valence (positive versus negative), and Temporal Distance (near versus far) of the imagined event, and parametric event modulators for the Rate Vividness and Rate Valence epochs (the participant’s rating). In our initial analyses, we modeled the entire 12 second Imagine epoch with one regressor/modulator. Then, to further examine the temporal order of vividness and valence effects, we modeled the first 4 seconds, middle 4 seconds, and last 4 seconds of the Imagine epoch with separate regressors/modulators.

Group analyses focused on the 12s imagination epoch. The main goal of this study was to examine dissociable roles of known DMN subcomponents (identified from previous literature) in future imagination. We first conducted region of interest analyses (ROIs) with masks from Shirer et al. (2012) for the dorsal and ventral default mode networks, since full maps of these networks were available for download. Shirer et al. (2012) applied FSL’s MELODIC independent component analysis (ICA) software at the group level for 15 participants who had completed a resting state scan and defined 90 ROIs. These results were confirmed using peak coordinates for the core and medial temporal lobe (MTL) components of the DMN obtained from Andrews-Hanna et al. (2010). Andrews-Hanna et al. (2010) used seed-based functional connectivity procedures to define regions that comprised each subsystem. The mean activity for each DMN subsystem was extracted from the specified image mask using the fslstats command. While the two ways of dividing the DMN are not identical, there is substantial overlap between the dorsal DMN and the DMN core, and between the ventral DMN and the DMN MTL subsystem.

Whole-brain group-level analyses were also performed to assess statistical significance at the whole-brain level. Corrected p-values were determined using permutation testing on the basis of cluster mass, with the cluster-defining threshold set to the p < 0.001 level (FSL *randomise*; 5000 iterations), and results were thresholded at corrected p < 0.05 to account for multiple comparisons.

## Results

### Behavioral Results

Behavioral ratings confirmed that we had successfully manipulated the vividness and valence of imagined events (Figure 2). There was a significant effect of vividness (*F*(3,45) = 31.54, *p* < 0.001) as well as valence (*F*(3,45) = 553.91, p < 0.001) in a one-way ANOVA across the four conditions. Vividness ratings were significantly different between vivid (mean = 5.32, SD = 0.36) and non-vivid (mean = 4.39, SD = 0.41) scenarios (*t*(31) = 10.34, p < 0.01), but not between positive (mean = 4.9, SD = 0.59) and negative (mean = 4.8, SD = 0.63) scenarios (*t*(31) = 0.84, p = 0.42). Valence ratings were significantly different between positive (mean = 5.72, SD = 0.34) and negative (mean = 2.14, SD = 0.35) scenarios (*t*(31) = 41.15, p < 0.01). Valence ratings were also slightly more positive for vivid (mean = 4.04, SD = 1.90) than non-vivid (mean = 3.82, SD = 1.80) scenarios (*t*(31) = 2.29, p = 0.03). Note that each scenario was presented twice, once with a “in the near future” prompt and once with a “in the far future” prompt, but there were no behavioral effects of near versus far future.

**Figure 2.**
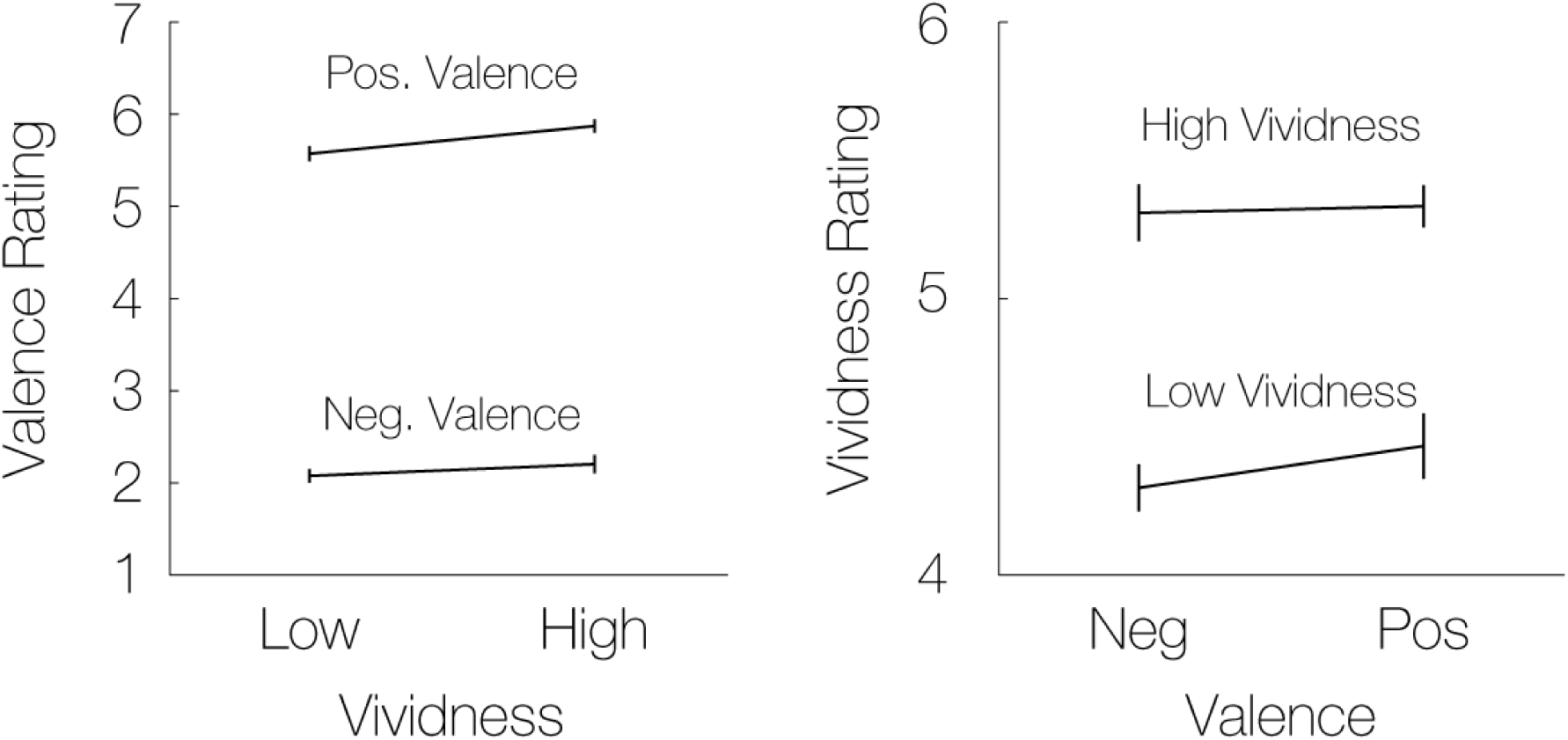
Behavioral manipulation check. The average valence and vividness ratings are shown four each of the four conditions (2 × 2 manipulation of vividness and valence). Left panel shows that the valence ratings are high for positive valence conditions and low for negative valence conditions, with little difference across vividness conditions. Right panel shows that the vividness ratings are high for high vividness conditions and low for low vividness conditions, with little difference across valence conditions.

### Imaging Results

The vividness and valence of imagined events modulated activity in distinct parts of the DMN. To test our hypothesis regarding differential functional roles of previously defined DMN subcomponents, we first examined the division of the DMN into dorsal and ventral components, as described by Shirer et al. (2012). The dorsal DMN was significantly modulated by valence (mean = 3.65, SE = 1.35) (*t*(23) = 2.71, *p* = 0.01), but not vividness (mean = −1.14, SE = 1.26) (*t*(23) = −0.90, *p* = 0.38), and the effect of valence was significantly larger than that of vividness (mean = −4.79, SE = 1.63) (*t*(23) = −2.95, *p* = 0.007). The ventral DMN was significantly modulated by vividness (mean = 3.65, SE = 1.07) (*t*(23) = 3.41, *p* = 0.002), but not by valence (mean = −0.63, SE = 0.98) (*t*(23) = −0.65, *p* = 0.52), and the effect of vividness was significantly larger than that of valence (mean = 4.28, SE = 1.36) (*t*(23) = 3.14, *p* = 0.005) (Figure 3). In addition, valence modulated the dorsal DMN significantly more than the ventral (*t*(23) = −4.48, *p* < 0.01), while vividness modulated the ventral DMN significantly more than the dorsal (*t*(23) = 5.51, *p* < 0.01).

**Figure 3.**
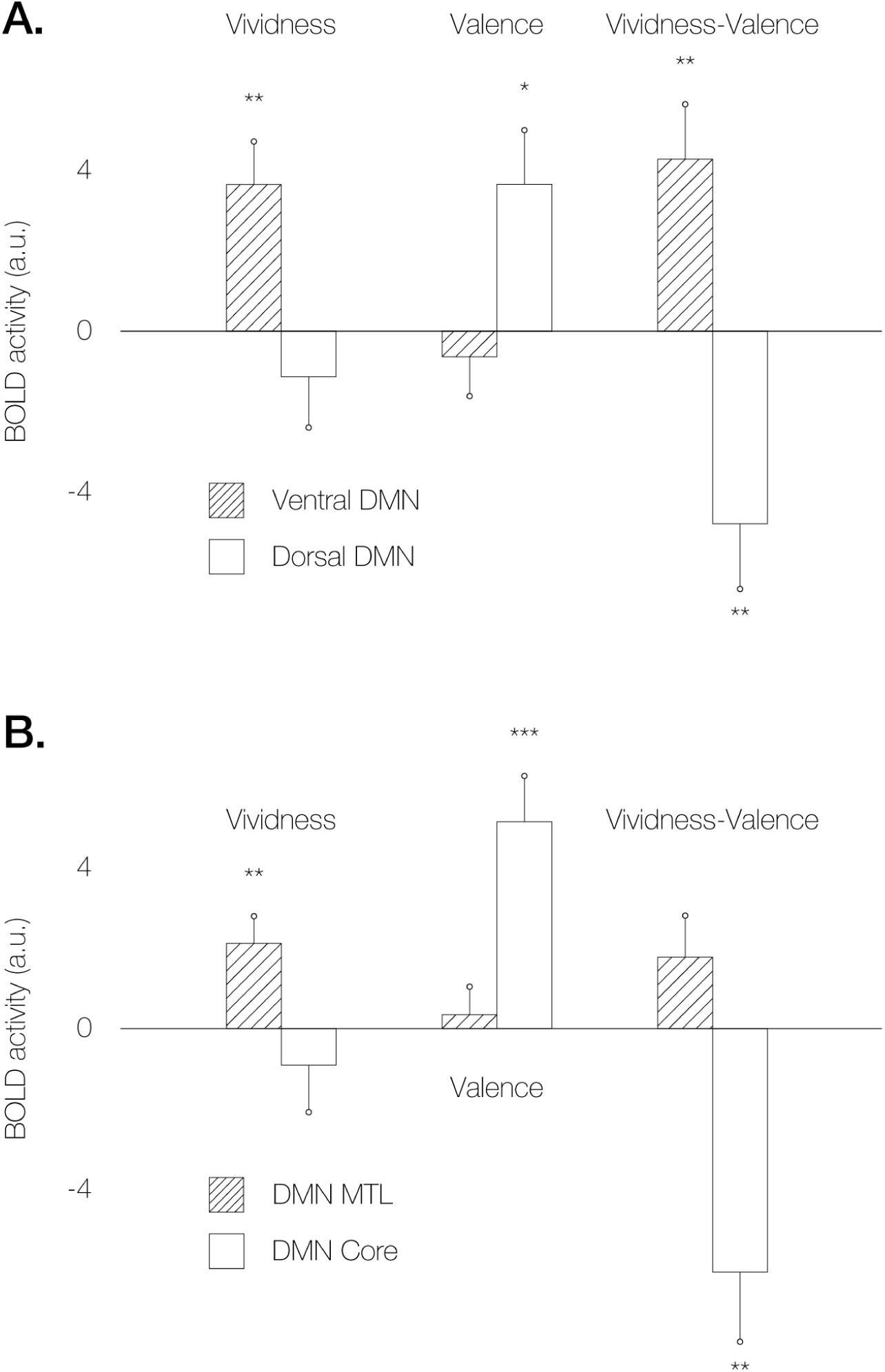
Panel A shows ROI results from the ventral and dorsal DMN, demonstrating that vividness but not valence significantly modulates the ventral DMN, while valence but not vividness significantly modulates the dorsal DMN. Panel B shows ROI results from the DMN medial temporal lobe (MTL) subregions and DMN core, demonstrating that vividness but not valence significantly modulates the DMN MTL system, while valence but not vividness significantly modulates the DMN core (* *p* <. 05, ** *p* < 0.01, *** *p* <. 001).

The dorsal and ventral DMN identified by Shirer et al. (2012) substantially overlap with the DMN core and DMN MTL divisions as described by Andrews-Hanna and colleagues (2007, 2010), and we observed the same dissociation using the peak coordinates from their study. The DMN core was significantly modulated by valence (*t*(23) = 4.52, *p* < 0.01), but not vividness (*t*(23) = −0.78, *p* = 0.44), and the effect of valence was significantly larger than that of vividness (*t*(23) = −3.45, *p* < 0.01). The MTL subsystem of the DMN was significantly modulated by vividness (*t*(23) = 3.13, *p* < 0.01), but not valence (*t*(23) = 0.49, *p* = 0.63), and the effect of vividness over valence was marginally significant (*t*(23)) = 1.75, *p* = 0.09) (Figure 3). In addition, valence modulated the DMN core significantly more than the DMN MTL (*t*(23) = 6.54, *p* < 0.01), while vividness modulated the DMN MTL significantly more than the DMN core (*t*(23) = −3.84, *p* < 0.01). Note that Andrews-Hanna and colleagues also described a dorsomedial DMN subsystem, but these regions exhibited no significant effects of either vividness (*t*(23) = – 1.62, *p* = 0.11) or valence (*t*(23) = −0.62, *p* = 0.54).

Whole-brain analyses further confirmed the separate modulation of distinct neural regions by vividness and valence (Figure 4). Across the entire imagination period, for trials with high compared to low vividness, there was increased activity in the left hippocampus, left dorsolateral prefrontal cortex (dlPFC), and bilateral orbitofrontal cortex (OFC). For trials with positive compared to negative valence, there was increased activity in the vmPFC and striatum. Furthermore, activation in the vmPFC was modulated more by valence than by vividness.

**Figure 4.**
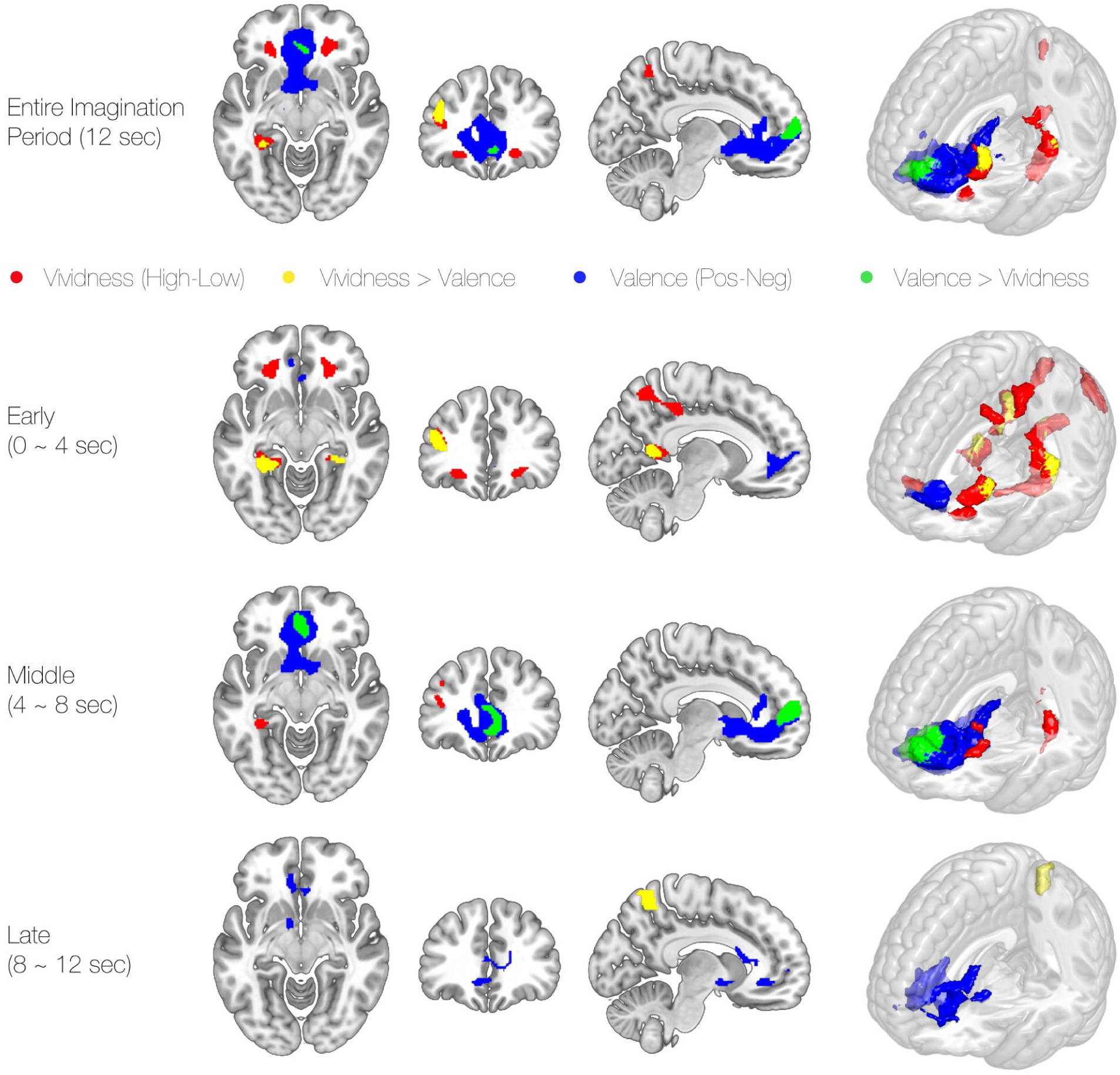
Whole-brain analysis of vividness and valence. Top panel shows the main effect of valence and vividness as well as their difference contrasts for the entire 12 second imagination period. The bottom three panels show the four effects for the early (first 4 s), middle (middle 4 s), and late (last 4 s) parts of the imagination period. List of the regions with their MNI coordinates are provided in Table 2.

Consistent with a temporal sequence of construction followed evaluation of imagined events, the effects of vividness were earlier in the imagination period than the effects of valence. We divided the imagination period into early (first 4s), middle (middle 4s), and late (last 4s) epochs. In the early epoch, the effects of vividness are most evident, with more vivid scenarios eliciting greater activity in the left dlPFC, bilateral hippocampus, retrosplenial cortex, precuneus and bilateral OFC. Activity in the bilateral hippocampus, left dlPFC, and right restrosplenial cortex were also modulated significantly more by vividness than by valence. In the middle epoch, effects of valence are most evident, with positive scenarios eliciting greater activation of the vmPFC and striatum and vmPFC being modulated significantly more by valence than by vividness. In the late epoch, the effects of vividness and valence have mostly subsided, though effects of valence persist in vmPFC and effects of vividness persist in precuneus (Table 2).

**Table 2.** Regions significantly modulated by vividness or valence in the whole-brain analyses at different time points. Regions such as bilateral hippocampus, bilateral OFC, left dlPFC, left precuneus, and right retrosplenial cortex were modulated by vividness, while vmPFC and VS were modulated by valence of imagined events.

| Description | Epoch | X | Y | Z |
| --- | --- | --- | --- | --- |
| *Vividness (High > Low)* |  |  |  |  |
| L Hippocampus | Entire period (0~12s) | -30 | -40 | -4 |
|  | Early (0~4s) | -30 | -36 | -14 |
|  | Middle (4~8s) | -30 | -40 | -4 |
| R Hippocampus | Early (0~4s) | 24 | -36 | -14 |
| L OFC | Entire period (0~12s) | -22 | 32 | -10 |
|  | Early (0~4s) | -24 | 36 | -10 |
| R OFC | Entire period (0~12s) | 22 | 34 | -10 |
|  | Early (0~4s) | 20 | 32 | -10 |
| L DLPFC | Entire period (0~12s) | -38 | 38 | 14 |
|  | Early (0~4s) | -40 | 40 | 14 |
|  | Middle (4~8s) | -38 | 40 | 14 |
| L Precuneus | Entire period (0~12s) | -10 | -58 | 56 |
|  | Early (0~4s) | -12 | -52 | 50 |
| L Occipital Mid. | Early (0~4s) | -30 | -82 | 46 |
| L Frontal Sup. | Early (0~4s) | -20 | 10 | 58 |
| R Retrosplenial/PCC | Early (0~4s) | 24 | -56 | 18 |
| *Vividness > Valence* |  |  |  |  |
| L Hippocampus | Entire period (0~12s) | -32 | -44 | -4 |
|  | Early (0~4s) | -32 | -38 | -12 |
| R Hippocampus | Early (0~4s) | 26 | -36 | -12 |
| L DLPFC | Entire period (0~12s) | -42 | 34 | 22 |
|  | Early (0~4s) | -40 | 38 | 14 |
| R Retrosplenial/PCC | Early (0~4s) | 14 | -50 | 12 |
| L Precuneus | Late (8~12s) | -12 | -54 | 56 |
| *Valence (Pos > Neg)* |  |  |  |  |
| vmPFC/VS | Entire period (0~12s) | -10 | 58 | 6 |
|  | Early (0~4s) | -2 | 42 | 4 |
|  | Middle (4~8s) | -10 | 58 | 6 |
|  | Late (8~12s) | 4 | 28 | -6 |
| *Valence > Vividness* |  |  |  |  |
| vmPFC | Entire period (0~12s) | -8 | 60 | 10 |
|  | Middle (4~8s) | -6 | 60 | 10 |

Though participants imagined each scenario twice, once in the near future and once in the far future, we did not observe any significant differences in activity between these two prompts, in either whole-brain or ROI analyses.

## Discussion

Our results demonstrate a functional double dissociation within the DMN by showing the separate modifiability of different subcomponents of the DMN by different aspects of imagination. We manipulated the vividness of imagined events to engage constructive processes during imagination and the valence of imagined events to engage evaluative processes. Vividness, but not valence, modulated activity in the ventral DMN, or DMN MTL subsystem, including precuneus and medial temporal lobe. Valence, but not vividness, modulated activity in the dorsal DMN, or DMN core, including the vmPFC. This basic pattern held in region of interest analyses using two different sets of DMN ROIs, as well as in whole-brain analyses. Vividness-modulated activity also happened early in the imagination period, while valence-modulated activity followed later in the imagination period. These findings support functional specialization within the DMN, with the ventral DMN/MTL subsystem involved in the construction of imagined future events and the dorsal DMN/core involved in the evaluation of imagined future events.

As proposed by Sternberg (2001), the kind of separate modifiability demonstrated here between the dorsal and ventral DMN provides strong evidence for dissociable mental modules. The logic of separate modifiability is similar to that of the canonical double dissociation, though importantly focuses on dissociations between processes within the context of a single task, rather than on dissociations between tasks. Key to the inferential strength of separate modifiability is that different measures (in our case, neural activity in the dorsal and ventral DMN) are shown to be both sensitive and specific (i.e., responding to some manipulations but not others). Given the demonstration of separate modifiability, we can infer that the single complex process of imagination can be decomposed into component processes—putatively, construction and evaluation—which can each be uniquely influenced by the distinct factors of vividness and valence.

Our findings complement and expand on prior work regarding the role of the DMN in imagination. Many previous studies have shown that imagination and other forms of “mental time travel” engage the DMN as a whole (Botzung et al., 2008; Hassabis et al., 2007; Hassabis & Maguire, 2007; Schacter et al., 2007). The DMN has greater metabolic activity at “rest”, when participants are left undisturbed to generate spontaneous thought, than during different executive cognitive tasks (Andrews-Hanna et al., 2010, 2007; Antoine Bechara, Damasio, Damasio, & Anderson, 2013; Fox, Spreng, Ellamil, Andrews-Hanna, & Christoff, 2015; Greicius et al., 2004; Raichle, 2015; Raichle et al., 2001; Shulman et al., 1997; Spreng et al., 2009). The DMN is also reliably activated when people engage in mental time travel in other ways, such as during tasks demanding autobiographic and social cognition, when people recall themselves in the past or think about someone else’s mental perspective (Addis et al., 2007; Atance & O’Neill, 2001; D’Argembeau et al., 2014, 2008; Schacter et al., 2007; Sharot et al., 2007; Spreng, Gerlach, Turner, & Schacter, 2015; Spreng et al., 2009; Tamir, Bricker, Dodell-Feder, & Mitchell, 2015). With respect to valence, several previous studies have shown, as we do here, that vmPFC is more active when participants imagine positive compared to negative scenarios (D’Argembeau & Van Der Linden, 2004; Gilbert & Wilson, 2007; Sharot et al., 2007). Though no previous studies have directly manipulated the vividness of imagined events, hippocampus and precuneus, the regions we find are more active when participants imagine more vivid scenarios compared to less vivid scenarios, are known to be important in episodic scene construction (Addis et al., 2007; Andrews-Hanna et al., 2010; Fletcher et al., 1995; Hassabis et al., 2007; Hassabis & Maguire, 2007; Schacter et al., 2007; Vincent et al., 2006).

Andrews-Hanna and colleagues (2010) first proposed that different sub-divisions of the DMN serve different constructive and evaluative functions, with what they called the MTL subsystem of the DMN (including hippocampus and precuneus) involved in the construction of mental scenes based on memory, and what they called the core DMN (including the vmPFC) involved in the affective evaluation of personal significance. The evidence to support this claim, though, was a single (as opposed to double) dissociation in which the MTL subsystem was more active when thinking about future than present events, while the DMN core was equally active in both conditions. However, other studies have observed different patterns of activity for thinking about the future versus the present, suggesting an alternative subdivision of the DMN into anterior and posterior components (Xu, Yuan, & Lei, 2016). Furthermore, subsequent studies of resting-state functional connectivity (Sestieri, Corbetta, Romani, & Shulman, 2011; Uddin, Kelly, Biswal, Castellanos, & Milham, 2009; Xu et al., 2016), meta-analytic co-activation (Laird et al., 2009, 2013), or task fMRI dissociations have yielded yet additional proposals regarding DMN sub-divisions (Bado et al., 2014; Leech, Kamourieh, Beckmann, & Sharp, 2011; Sestieri et al., 2011; Whitfield-Gabrieli et al., 2011; Xu et al., 2016). Therefore, the separate modifiability by vividness and valence demonstrated here provides much stronger support for the distinction originally proposed by Andrews-Hanna and colleagues, between DMN components involved in constructive and evaluative processes during imagination.

Though we manipulated vividness to engage constructive processes and valence to engage evaluative processes, we do not expect that the activity modulations observed are necessarily unique to these specific features, as opposed to any set of features that would differentially engage construction versus evaluation. A broader set of potential features are seen in the more comprehensive ratings of our scenarios (Table 1). Vivid scenarios were also more likely to be imagined from a first person viewpoint, while positive scenarios were also higher in arousal, social and temporal connectedness, and self-relevance. Whether these features can be dissociated from each other, and whether some subset can be shown to be the primary driver of activity in the ventral or dorsal DMN, are important questions for future research. Regardless of the answer to these questions, the current results establish a clear dissociation between the roles of the ventral and dorsal DMN in constructive versus evaluative processes during imagination.

We also identified a few regions outside the DMN that were engaged by vividness or valence. The ventral striatum (VS) was more active when imagining positive compared to negative events. This is consistent with the involvement of the vmPFC and VS in evaluating outcomes and encoding predicted value during decision-making tasks (Bartra et al., 2013; Kable & Glimcher, 2007). Interestingly, we observed activation in bilateral orbitofrontal cortex (OFC) for more vivid compared to less vivid scenarios. Several lines of evidence implicate the OFC in decision-making as well, with recent theories proposing that the OFC represents specific outcomes, rather than value itself, that are necessary for computing value and planning during choice tasks (A. Bechara, 2000; Ursu & Carter, 2005; Wallis, 2007). Though there remains much debate over the extent to which OFC and vmPFC may play distinct roles in decision-making, our finding that vmPFC is specifically modulated by valence during imagination, while central OFC is specifically modulated by vividness, supports a distinction between these two closely related areas.

Recent studies suggest that humans spend most of their time engaged in mental time travel, either remembering the past or imagining the future (Killingsworth & Gilbert, 2010). Yet we have very little formal understanding of the psychological processes involved in imagination. Our results suggest that the complex process of imagination, which might appear to be unitary, can in fact be decomposed into (at least) two dissociable mental processes, the construction of novel potential future events from components in memory and the evaluation of constructed events as desirable or undesirable. Neural measurements provided the key evidence for this dissociation, given the difficulty in constructing objective behavioral measures of imagination quality or ability. Thus, neuroscientific methods may prove critical to the further understanding of this central aspect of human subjective experience.

The authors declare no competing interests

## Acknowledgements

This study was supported by National Institute of Drug Abuse grant R01 DA029149.

